# Urine proteome changes in a chronic unpredictable mild stress (CUMS) mouse model of major depressive disorder

**DOI:** 10.1101/2020.06.17.156265

**Authors:** Yuhang Huan, Jing Wei, Tong Su, Youhe Gao

**Affiliations:** Department of Biochemistry and Molecular Biology, Beijing Normal University, Gene Engineering Drug and Biotechnology Beijing Key Laboratory, Beijing, China; State Key Laboratory of Medical Molecular Biology, Institute of Basic Medical Sciences Chinese Academy of Medical Sciences and Peking Union Medical College, Neuroscience Center, Chinese Academy of Medical Sciences, Beijing, China

**Keywords:** Major depressive disorder, CUMS model, urine proteome, biomarker

## Abstract

**Background:** Major depressive disorder (MDD) is a prevalent complex psychiatric disorder with a high prevalence rate. Because MDD is a systemic multifactorial disorder involving complex interactions and disturbances of various molecular pathways, there are no effective biomarkers for clinical diagnosis. Urine is not subjected to homeostatic control, allowing it to reflect the sensitive and comprehensive changes that occur in various diseases. In this study, we examined the urine proteome changes in a CUMS mouse model of MDD.

**Methods:** Male C57BL/6 mice were subjected to chronic unpredictable mild stress for 5 weeks. The tail suspension test (TST) and sucrose consumption test (SCT) were then applied to evaluate depression-like behaviors. The urine proteomes on day 0 and day 36 in the CUMS group were profiled by liquid chromatography coupled with tandem mass spectrometry (LC-MS/MS).

**Results:** A total of 45 differential proteins were identified, 24 of which have been associated with the pathogenic mechanisms of MDD, while 10 proteins have been previously suggested as MDD biomarkers. There was an average of two differential proteins that were identified through 1048574 random combination statistical analyses, indicating that at least 95% of the differential proteins were reliable and not the result of random combination. The differential proteins were mainly associated with blood coagulation, inflammatory responses and central nervous system development.

**Conclusions:** Our preliminary results indicated that the urine proteome can reflect changes associated with MDD in the CUMS model, which provides potential clues for the diagnosis of clinical MDD patients.

## Introduction

Major depressive disorder (MDD) is a serious mental illness affecting 15% of individuals throughout their lifetime [1]. The World Health Organization (WHO) claims that MDD will be the leading cause of burden due to disability and diseases globally by 2030. Usually, the clinical diagnosis of MDD is based on patient interviews, checklists and questionnaires [2]. In this context, the identification of candidate biomarkers associated with MDD is crucial. Various studies have aimed to identify candidate MDD biomarkers [3–5]. However, as MDD is a multifactorial and highly heterogeneous disease, it is still challenging to identify effective biomarkers for MDD clinical diagnosis.

As the filtration of blood, urine is not regulated by the homeostatic mechanism of the body. Therefore, earlier and more subtle changes associated with physiological and pathological changes may be reflected in urine [6]. Urine proteomics has already been applied to identify potential MDD biomarkers [7–9]. However, most studies used clinical urine samples from MDD patients, which are easily affected by various influential factors, such as age, gender, diet and some necessary medications. Using animal models is an effective way to determine the direct relationships of biomarkers with diseases, as the use of models can minimize the influence of such external factors [10]. In our previous studies, early candidate urine biomarkers could be detected successfully before ±-amyloid deposition in a APP/PS1 transgenic mouse model of Alzheimer’s disease (AD) [11]. In addition, several urinary differential proteins were screened in an a-synuclein transgenic mouse model of Parkinson’s disease (PD) [12]. Since recent evidence suggests that there is a strong relationship between MDD and neurodegenerative diseases such as AD and PD [13, 14], we therefore aimed to determine whether the urine proteome can reflect changes associated with MDD.

The chronic unpredictable mild stress (CUMS) model is an animal model commonly used to simulate the core behavioral characteristics of human depression [15, 16]. Therefore, the CUMS model is suitable for investigating the pathophysiology of depression and assisting in diagnosis. In this study, the CUMS mouse model was established by exposing mice to mild stressors in a random order for 35 days. Then, all of the CUMS mice were evaluated by behavioral tests, such as the tail suspension test (TST) and sucrose consumption test (SCT). In order to contain all the effectors of stressors, we collected urine samples from the CUMS mouse models on days 0 and 36. The urinary proteomes were profiled using liquid chromatography coupled with tandem mass spectrometry (LC-MS/MS) with the data independent acquisition (DIA) method. This study was designed to identify urine proteome changes associated with MDD. The technical flowchart is presented in Figure 1.

**Figure 1.**
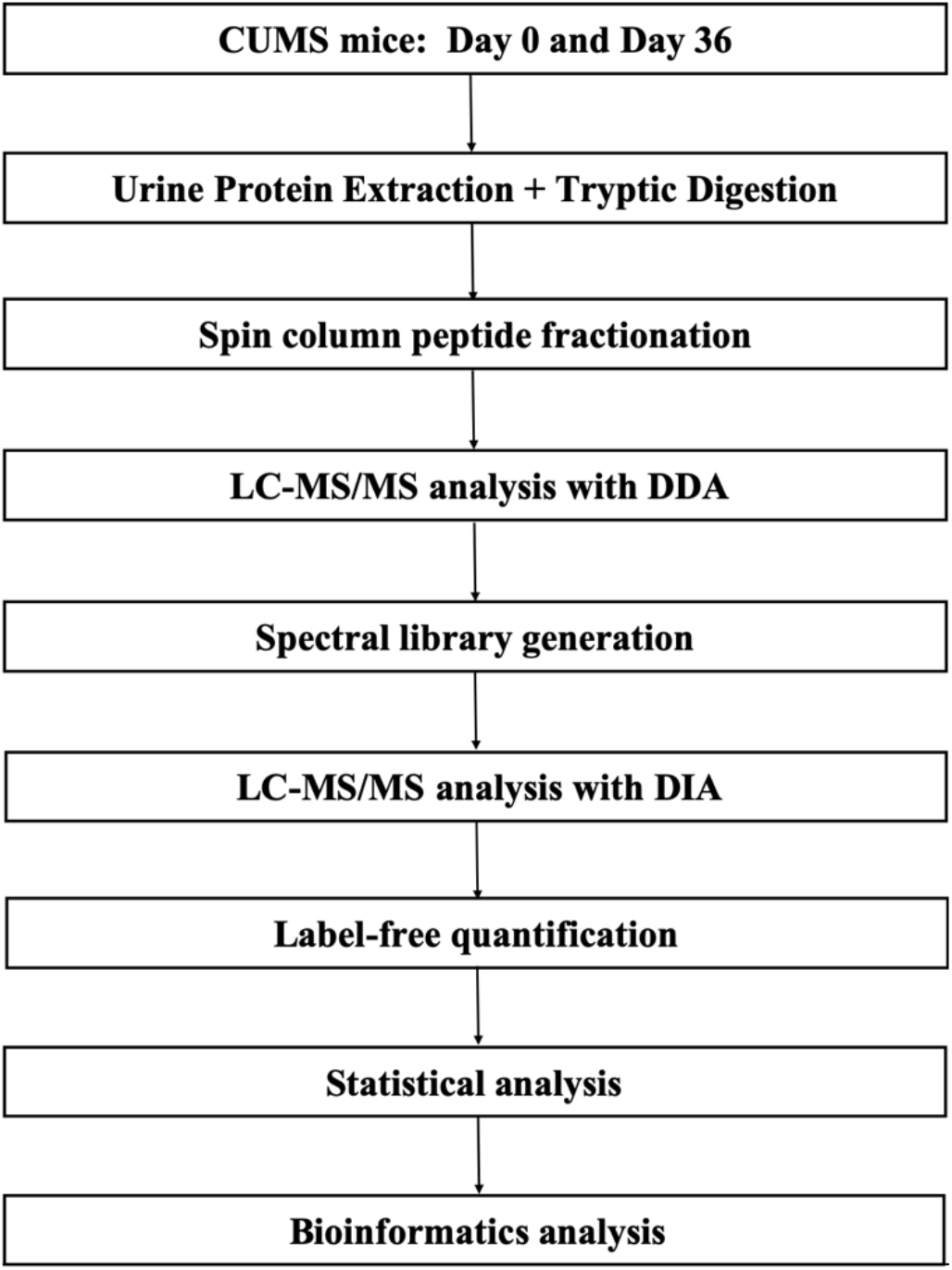
Workflow of protein identification in CUMS mice. Urine samples were collected from the CUMS mice on days 0 and 36. The urinary proteomes were profiled using liquid chromatography coupled with tandem mass spectrometry (LC-MS/MS). Bioinformatics analysis was performed using DAVID (https://david.ncifcrf.gov) and STRING (https://string-db.org/cgi/input.pl).

## Materials and methods

### The establishment of the CUMS model

All animal experiments were approved by the Ethics Review Committee of the Institute of Basic Medical Sciences, Chinese Academy of Medical Sciences. C57BL/6 male mice (n = 32, 8 weeks) were purchased from Beijing Vital River Laboratory. The mice were acclimated to the environment for one week before the experiment and kept under standard conditions with a 12-hour light/dark cycle, a temperature of 23±1°C, a humidity of 50%, and free access to standard rodent chow and water. All experimental animals were utilized in accordance with the “Guidelines for the Care and Use of Laboratory Animals” issued by the Beijing Office of Laboratory Animal Management (Animal Welfare Assurance Number: ACUC-A02-2015-004).

The CUMS model was established according to previously reported procedures [17, 18]. In brief, the C57BL/6 male mice were divided into the control group (n=20) and the CUMS group (n=12). Then, the CUMS group was exposed to a variety of mild stressors: (1) overnight light exposure, (2) darkness for 12 h, (3) wet bedding for 12 h, (4) removal of bedding for 12 h, (5) tilting of the cage (45°) for 12 h, (6) deprivation of food for 12 h, (7) deprivation of water for 12 h, (8) soaking for 3 min in cold water (4 °C), (9) soaking for 3 min in water at 45 °C, (10) physical restraint for 4 h, (11) exposure to the smell of cats for 3 h, (12) exposure to the sound of cats for 3 h, (13) cage vibration (80 rpm) for 3 h, (14) strobe lighting for 3 h, (15) and electrical shocks to the feet for 15 min (0.2 mA). The CUMS group was randomly subjected to the various stressors repeatedly for a period of five weeks. A combination of three different stressors was applied each day, and the same stressor was not repeated within three days.

### Behavioral tests

The tail suspension test (TST) was performed at baseline and after the CUMS procedure (Day 38) to assess the state of despair. The mice were suspended by their tails from an acrylic rod (15 cm in diameter, 30 cm in height) for six min and monitored using a video tracking system (Smart 3.0). Struggle-related behavior was assessed, and the immobility time during the 6 min suspension period was recorded.

The sucrose consumption test (SCT) was used to assess anhedonia in mice. The SCT was performed at baseline and on days 39-42. First, we acclimated the animal to 1% sucrose solution using a homemade water bottle (50 ml centrifuge tube with plastic plugs) for two days and then caged each animal individually without water for 12 h on the third night. Finally, on the morning of the fourth day, the mass of the 1% sucrose solution was measured by an electronic balance after allowing the mice to drink for 1 h. The consumption of 1% sucrose solution was calculated. The details of the time processing of the behavioral tests are presented in Figure 2.

**Figure 2.**
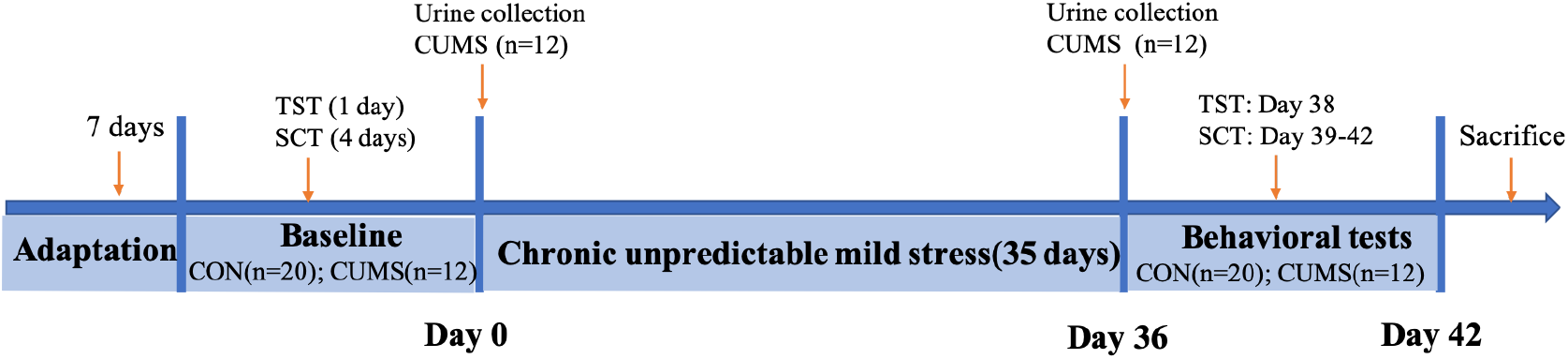
CUMS modeling and the behavioral testing process. C57BL / 6 male mice were divided into the control group (n=20) and CUMS group (n=12). The CUMS group was randomly subjected to the various stressors repeatedly for five weeks. The tail suspension test and sucrose consumption test were used to assess the effect on depression in the CUMS and the control group.

### Urine collection and sample preparation

Urine samples from twelve CUMS mice were collected in metabolic cages on days 0 and 36. Before the first urine sample collection, all mice were accommodated in metabolic cages for 3 days. All mice were placed in metabolic cages individually for 12 h to collect urine without any treatment (at least 1 mL). After collection, the urine samples were centrifuged at 3,000 × *g* for 30 min at 4°C and then stored at −80°C. For urinary protein extraction, the urine samples were first centrifuged at 12,000 g for 30 min at 4°C. Then, 500 μL urine from each sample was precipitated with three volumes of ethanol at −20°C overnight. The pellets were dissolved in lysis buffer (8 mol/L urea, 2 mol/L thiourea, 50 mmol/L Tris, and 25 mmol/L DTT). Finally, the supernatants were quantified by Bradford assay.

A total of 100 μg protein was digested with trypsin (Trypsin Gold, Mass Spec Grade, Promega, Fitchburg, WI, USA) using filter-aided sample preparation methods [19]. Briefly, the protein in each sample was loaded into a 10 kDa filter device (Pall, Port Washington, NY, USA). After washing two times with urea buffer (UA, 8 mol/L urea, 0.1 mol/L Tris-HCl, pH 8.5) and 25 mmol/L NH4HCO3 solutions, the protein samples were reduced with 20 mmol/L dithiothreitol (DTT, Sigma) at 37 °C for 1 h and alkylated with 50 mmol/L iodoacetamide (IAA, Sigma) for 45 min in the dark. The samples were then washed with UA and NH4HCO3 and digested with trypsin overnight at 37°C. The digested peptides were desalted using Oasis HLB cartridges (Waters, Milford, MA, USA) and then dried by vacuum evaporation (Thermo Fisher Scientific, Bremen, Germany).

### Spin column peptide fractionation

The digested peptides were dissolved in 0.1% formic acid and diluted to a concentration of 0.5 μg/μL. To generate the spectral library, a pooled sample (96 μg, 4 μg of each sample) from 24 samples was loaded onto an equilibrated, high-pH, reversed-phase fractionation spin column (84868, Thermo Fisher Scientific). A step gradient of 8 increasing acetonitrile concentrations (5, 7.5, 10, 12.5, 15, 17.5, 20 and 50% acetonitrile) in a volatile high-pH elution solution was then added to the columns to elute the peptides as eight different gradient fractions. The fractionated samples were then evaporated using vacuum evaporation and resuspended in 20 μL of 0.1% formic acid. Two microliters of each fraction was loaded for the LC-DDA-MS/MS analysis.

### LC-MS/MS analysis

The iRT reagent (Biognosys, Switzerland) was added at a ratio of 1:20 v/v to all peptide samples to calibrate the retention time of the extracted peptide peaks. For analysis, 1 μg of peptide from each sample was loaded into a trap column (75 μm * 2 cm, 3 μm, C18, 100 Å) at a flow rate of 0.25 μL/min and then separated with a reversed-phase analytical column (75 μm * 250 mm, 2 μm, C18, 100 Å). Peptides were eluted with a gradient of 4%-35% buffer B (0.1% formic acid in 80% acetonitrile) for 90 min and then analyzed with an Orbitrap Fusion Lumos Tribrid Mass Spectrometer (Thermo Fisher Scientific, Waltham, MA, USA). The LC settings were the same for both the DDA-MS and DIA-MS modes to maintain a stable retention time.

For the generation of the spectral library, the ten fractions obtained from the spin column separation were analyzed with mass spectrometry in DDA mode. The MS data were acquired in high-sensitivity mode. A full MS scan was acquired within a 350-1,500 m/z range with the resolution set to 120,000. The MS/MS scan was acquired in Orbitrap mode with a resolution of 30,000. The HCD collision energy was set to 30%. The AGC target was set to 5e4, and the maximum injection time was 45 ms.

Twenty-four individual samples were analyzed in DIA-MS mode. The variable isolation window of the DIA method with 39 windows was used for DIA acquisition (Table S1). The full scan was obtained at a resolution of 60,000 with an m/z range from 350 to 1,400, and the DIA scan was obtained at a resolution of 30,000. The AGC target was 1e6, and the maximum injection time was 50 ms. The HCD collision energy was set to 32%. A single DIA analysis of pooled peptides was performed as a quality control after every 4th sample. Finally, a total of 6 QC samples were analyzed.

### Label-free DIA quantification

To generate a spectral library, ten DDA raw files were first searched by Proteome Discoverer (version 2.1; Thermo Scientific) with SEQUEST HT against the SwissProt Mus musculus database (released in May 2019, containing 17,038 sequences). The iRT sequence was also added to the *Mus musculus* database. The precursor and product ion spectra were searched at an initial mass tolerance of 10 ppm and 0.02 Da, respectively. Tryptic digestion was selected, and up to two missed cleavages were allowed. The dynamic modification was set as the oxidation (+15.995 Da) of methionine, and the static modification was the carbamidomethylation (+57.021 Da) of cysteine. The false discovery rate (FDR) of both proteins and peptides was set to 1%. Finally, a total of 647 protein groups from the library were identified. Then, the pdResult file was imported into Spectronaut™ Pulsar X (Biognosys, Switzerland) software to generate a spectral library [20].

The 6 QC and 24 raw DIA files were imported into Spectronaut Pulsar X with the default settings. The peptide retention time was calibrated according to the iRT data. Cross-run normalization was performed to calibrate the systematic variance of the LC-MS performance, and local normalization based on local regression was used [21]. Protein inference to generate the protein groups, was performed using the implemented IDPicker algorithm [22]. All results were then filtered according to a Q value less than 0.01 (corresponding to an FDR of 1%). The peptide intensity was calculated by summing the peak areas of the respective fragment ions for MS2. The protein intensity was calculated by summing the respective peptide intensity.

### Statistical analysis

The missing of abundance values were determined with the KNN method [23]. For later analysis, we required the proteins to be removed if the CVs of the QC samples were more than 30%. Comparisons between two groups were conducted using two-sided unpaired t-tests. The differential proteins were selected according to *P* <0.01 and a fold change ≥1.5 or ≤0.67.

### Bioinformatics analysis

The differential proteins were analyzed by Gene Ontology (GO) based on the biological process, cellular component, molecular function and pathway categories using DAVID [24]. Protein interaction network analysis was performed using STRING software (https://string-db.org/cgi/input.pl) based on the STRING database.

## Results

### Experimental results of depressive-like behavior in the CUMS model

The behavioral experiments were performed for the CUMS group (n=12) and the control group (n=20) to assess the despair state. The results of the tail suspension test (TST) are presented in Figure. 3A. There was no significant difference between the two groups before modeling, indicating that the mice were in a similar state at the beginning of the experiment. After 35 days of stress exposure, the immobility time of the CUMS group increased significantly compared with the initial value determined for the CUMS group before modeling (p<0.05) and the value determined for the control group (p<0.01). The SCT results are presented in Figure 3B. Before modeling, there was no significant difference in the baseline. After modeling, the sucrose consumption in the CUMS group was decreased compared with that before modeling (p<0.01) and in the control group (p<0.0001). The TST and SCT experiments indicated that the CUMS model successfully simulated the depressive-like behavior.

**Figure 3.**
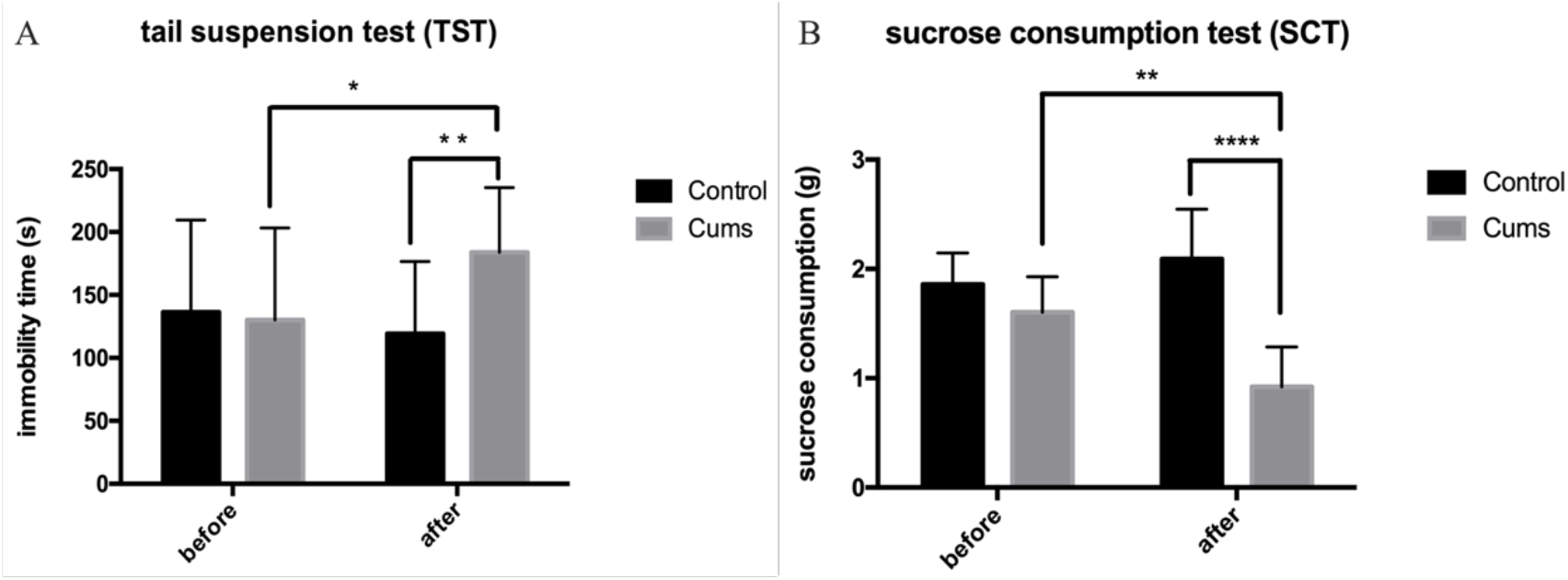
Experimental results of depressive-like behavior in the CUMS model. Twenty mice in the control group and twelve CUMS model mice were included in the experimental tests. (A). Results of the tail suspension test. (B). Results of the sucrose consumption test. All these results are presented as the mean ± SD. *p<0.05; **p<0.01; ****p<0.0001.

### Urine proteome changes in CUMS mice

A total of 522 protein groups were identified in the 24 biological replicates. Specifically, 266 proteins that were considered to be highly authentic were identified when the CV values of protein abundance for the QC samples and the missing values for all samples were both less than 30%. Based on a P value < 0.01 and a fold change ≥1.5 or ≤0.67, 45 differential proteins were selected. A hierarchical clustering of the 45 differential proteins was performed by using the average linkage method. As shown in Figure S1, all the differential proteins were clustered into two groups, which corresponded to the urine proteome changes found on day 0 and day 36, indicating that depression behavior in the CUMS model can be distinguished on the basis of these 45 differential proteins. A total of 39 differential proteins were homologous; 24 have been previously associated with the pathogenic mechanisms of MDD, while 10 have been suggested as MDD biomarkers previously. Additional details are presented in Table 1.

**Table1.**
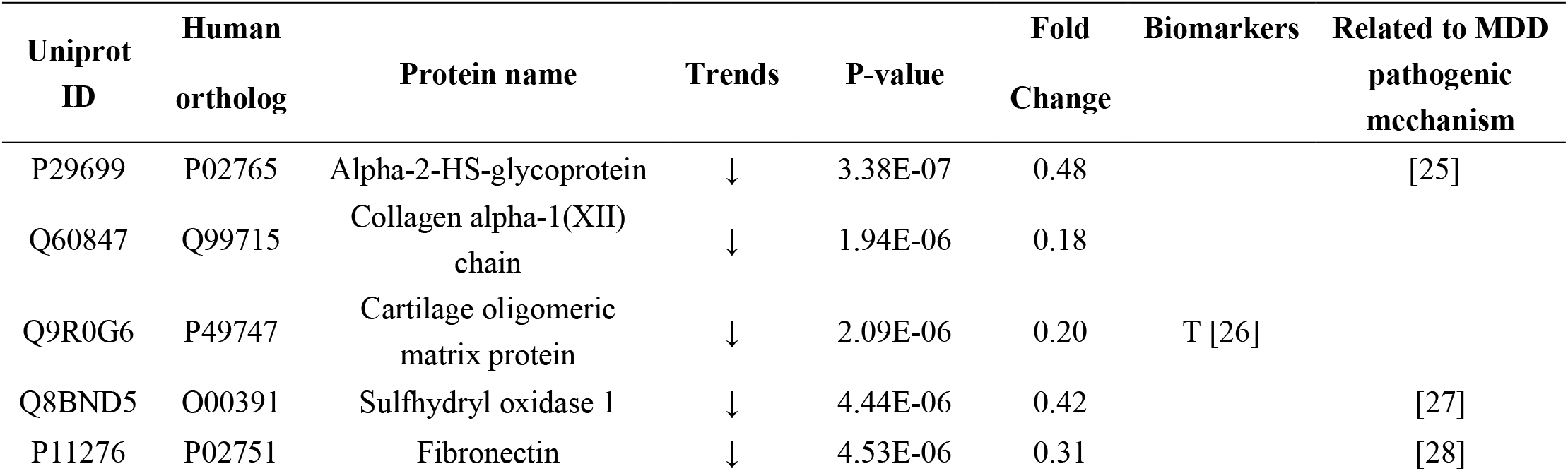
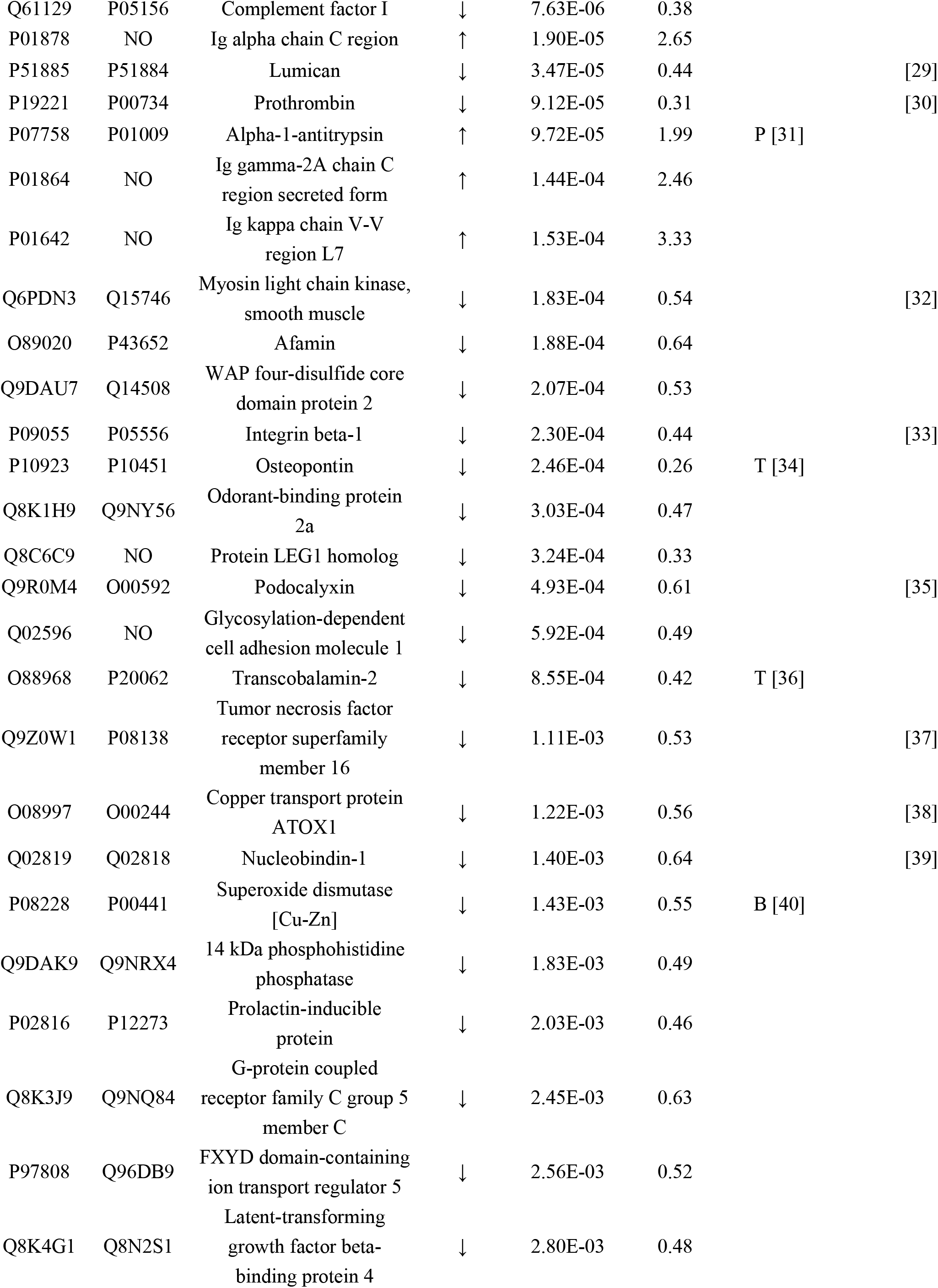
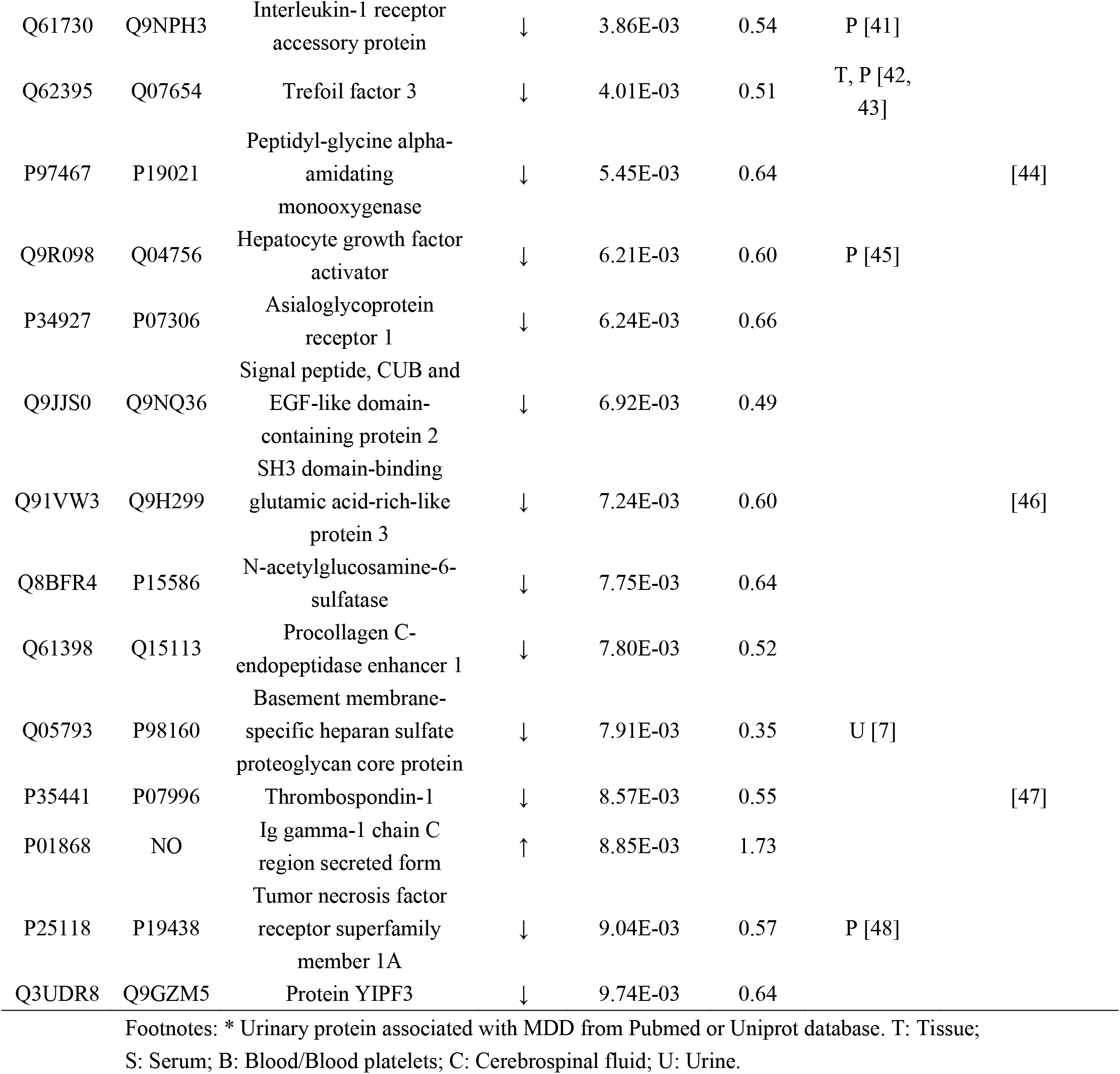
Differential proteins identified on day 0 and day 36.

The identified number of proteins in each sample is presented in Table S2. All the quantification details are listed in Table S3.

### Randomness generation rate of differential proteins

To determine the reliability rates of the 45 identified differential proteins, we randomly divided the 24 samples into two groups, and the numbers of differential proteins for each combination were generated. The screening criteria were the same as those used before. Finally, a total of 1048574 combinations were generated; all details are presented in Table S4. As a result, the average number of differential proteins in the 1048574 combinations was 1.64 (approximately two differential proteins), indicating that the 45 identified differential proteins could not be selected through the random combination statistical analysis. In addition, the percentage of randomly identified proteins was predicted to be 3.64%, indicating that at least 96.36% of the differential proteins were not identified randomly (Table 2). All these results indicated that the prediction of the 45 differential proteins was reliable, with little possibility of randomness.

**Table 2.** Results of the random combination statistical analysis.

| Screening criteria | Total number of random combinations | Average numbers of proteins with false random combinations | Percentage (average number of protein false random combinations/number of correctly identified differential proteins) |
| --- | --- | --- | --- |
| FC≥1.5 or ≤0.67<br>P-value <0.01 | 1048574 | 2 | 0.0364 |

### Functional annotation of the differential proteins

The functional enrichment analysis of the differential proteins was carried out using DAVID to examine the associated biological processes, cellular components, molecular functions and pathways. To obtain sufficient biological functional information, a significance threshold according to a P value <0.05 and a FC range ≥ 1.5 or ≤ 0.67 was applied. Finally, a total of 74 differential proteins were selected.

The list of the thirteen representative biological processes is presented in Figure 4A (all details are shown in Table S5). Acute-phase response, innate immune response, positive regulation of blood coagulation, inflammatory response, negative regulation of nitric oxide-mediated signal transduction, vitamin transport, negative regulation of neuron projection development and central nervous system development were enriched. For the cellular components, most of the differential proteins were associated with the extracellular space, extracellular region, extracellular exosome, extracellular matrix and blood microparticle (Fig 4B). For molecular functions, glycoprotein binding, protease binding, fibronectin binding and small molecule binding were overrepresented (Fig 4C). For the pathways, the ECM-receptor interaction, focal adhesion, proteoglycans in cancer, PI3K-Akt signaling, and complement and coagulation cascade pathways showed obvious changes (Fig 4D).

**Figure 4.**
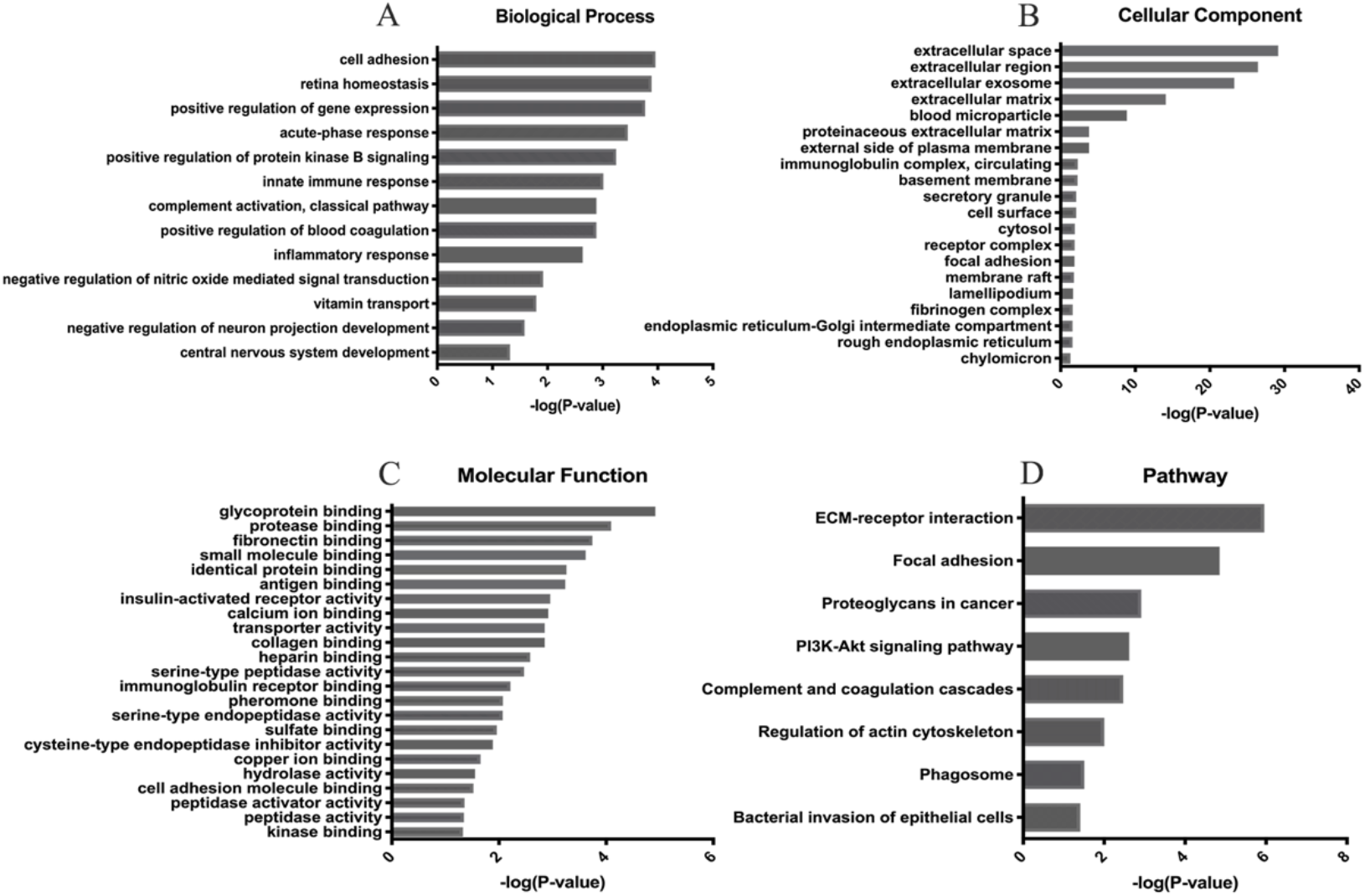
Functional annotation of the differential proteins. Functional annotations were generated by GO analysis. Lists of thirteen representative biological processes (A), cellular components (B), and molecular functions (C). Significant pathways were enriched based on the KEGG database analysis (D). The criteria of p<0.05 and a fold change ≥ 1.5 or ≤ 0.67 were applied for screening.

### Interaction analysis of the 45 differential proteins in the CUMS model

The STRING database was used to map the protein interaction network of the differential proteins. The results are shown in Figure 5. The p-value of PPI enrichment was less than 1.0e-16. A total of thirteen proteins (F2, AHSG, SERPINA1A, SPP1, NUCB1, QSOX1, THBS1, FN1, ITGB1, COMP, LUM, PCOLCE and COL12A1) were closely related, and they were associated with the integrin pathway, platelet degranulation, neuroprotection and the regulation of reactive oxygen species metabolic processes.

**Figure 5.**
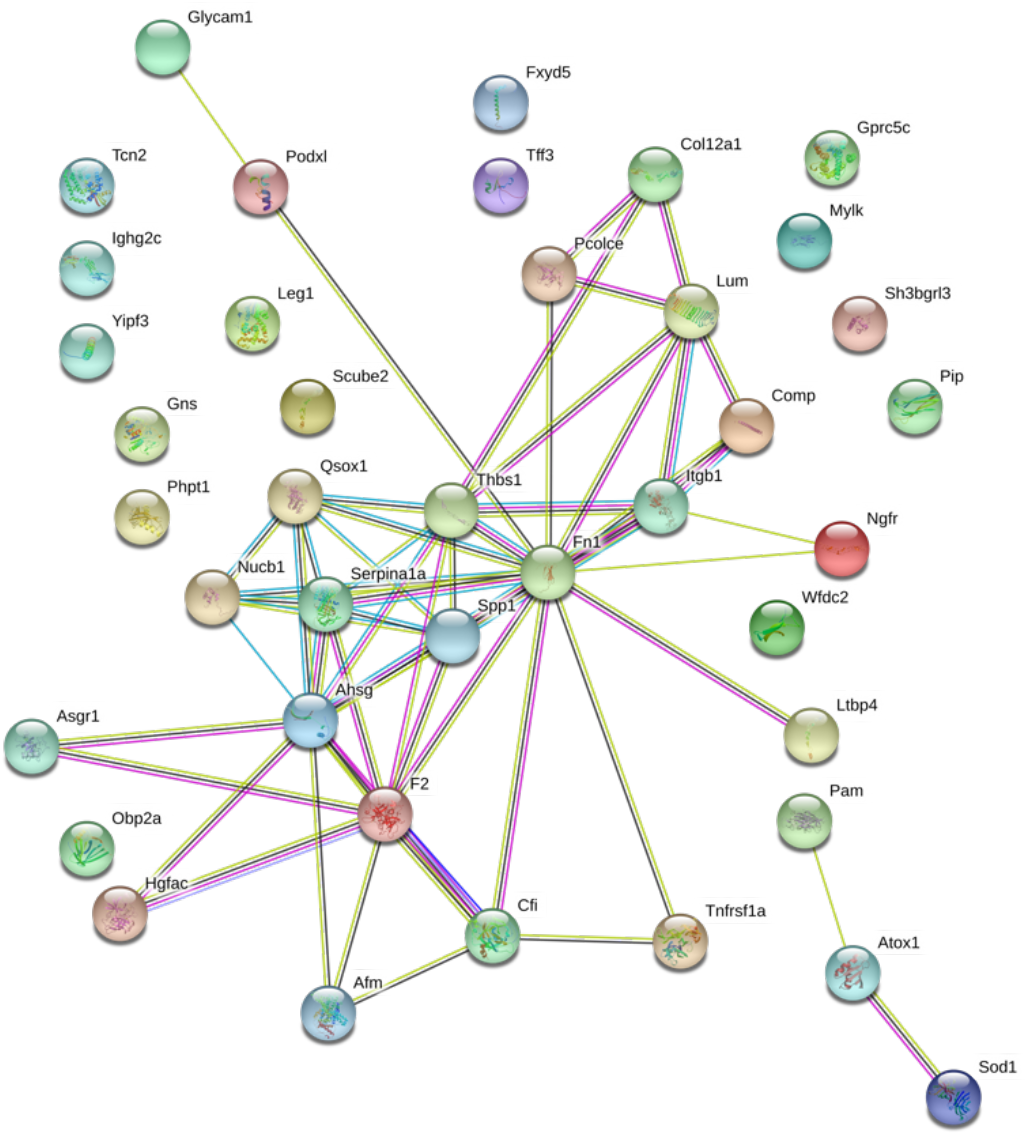
Interaction diagram of the 45 differential proteins. Forty-five differential proteins were involved in the protein interaction network based on the STRING database (p<0.01 and fold change ≥ 1.5 or ≤ 0.67).

## Discussion

After the functional enrichment analysis, some of the enriched biological processes were found to be associated with MDD. (i) For example, in association with the positive regulation of blood coagulation, increased reactivity and aggregation of platelets have been reported in major depression [49]. Both tricyclic antidepressants and selective serotonin reuptake inhibitors (SSRIs) act on serotonin transport receptor (SERT) to block the uptake of 5-HT by neurons and platelets [50]. Several differential proteins corresponding to the QSOX1, SERPINA1, COMP, F2 and THBS1 genes are involved in the process of blood coagulation and platelet degranulation. (ii) The acute phase response, inflammatory response and negative regulation of nitric oxide-mediated signal transduction were reported to play roles in MDD. The significantly changed biological process known as the acute phase response is a systemic response triggered by local inflammation. Inflammatory responses caused by long-term stress have been proven to be related to the occurrence of depression [51]. The F2, FN1, AHSG, SERPINA1, TNFRSF1A, IL1RAP, NGFR, SPP1, THBS1, MYLK, SPP1, IL1RAP and GPRC5B proteins are related to NF-κB, which mediates inflammatory pathway processes and NO production. NF-κB has been reported as an important therapeutic target for psychiatric disorders [52] and plays a key role in the regulation of synaptic signals and neuronal morphology [53]. The evidence shows that the increase in NO caused by chronic pressure can affect the activity of the HPA axis [54], which plays an important role in the development of depression [55]. (iii) NGFR, CLU, FN1 and PAM function in central nervous system development and may participate in the development of MDD.

Additionally, some pathways were also reported to play important roles in MDD: (i) According to the KEGG analysis, the BDNF-PI3K/Akt pathway was associated with the enriched proteins FN1, THBS1, ITGB1, NGFR and SPP1. The phosphoinositide 3-kinase (PI3K/Akt) pathway can reduce depression-like behaviors induced by neuroinflammation and plays an important role in the neuroprotective process of depression [56, 57]. Brain-derived neurotrophic factor (BDNF) functions as a growth factor and can activate the PI3K/Akt pathway in neurons. Regulation of BDNF may reverse stress-induced defects in the adult brain structure and synaptic plasticity and thus improve the ability to adapt/respond to environmental changes that may accelerate or exacerbate depressive episodes [58]. (ii) Proteins such as ITGB1, FN1, THBS1, COMP, HSPG2 and SPP1 participate in integrin family-related pathways. Integrins are a class of important cell surface receptors that mainly mediate adhesion between cells and ECM receptors. The binding process of the urokinase receptor and integrin mediates the processes of regulating cell adhesion, intercell signal transmission and axonal regeneration after injury in the central nervous system. [59]. Integrin-linked kinase (ILK) is also involved in the nerve growth factor-mediated postinjury repair of nerve fibers and binding in the PI3K/Akt pathway [60]. The results of the functional analysis showed that several MDD related biological processes and pathways involving a set of differential proteins can be identified through the urine proteome. Most likely, we should determine the panel used for the diagnosis of MDD by using a combination of differential proteins involved in several related processes.

Only one protein was identified by both our study and a quantitative plasma proteomic study performed in the CUMS model [61]. However, among the top five canonical pathways identified in the plasma study, three pathways were also be enriched in our study, including acute phase response signaling, complement system and intrinsic prothrombin activation. Although there was only one differential protein in common with the plasma proteomic study, the biological processes enriched by the two studies were similar. We therefore suggest that the urine proteome may also provide a possibility for MDD diagnosis.

We also compared the urinary differential proteins in the CUMS model to those in 15 other models, including a clinical model of autism [62] and a variety of animal models of Alzheimer’s disease [11], Parkinson’s disease [12], myocarditis [63], chronic pancreatitis [64], unilateral ureteral obstruction model [65], astrocytoma [66], liver fibrosis [67], pulmonary fibrosis [68], glomerulosclerosis [69], a model involving the injection of 10 tumor cells [70], the Walker-256 intracerebral tumor model [71], the Walker-256 liver tumor model [72], the Walker-256 subcutaneous model [73] and the Walker-256 lung metastasis model [74]. The comparison results are presented in Table S6. Interestingly, we found that there were eight differential proteins identified in both the CUMS and AD models, which was far more than that identified for other models. It has been reported that depression is a risk factor for the development of AD, and there are similar pathophysiological changes, such as the presence of hippocampal atrophy, oxidative stress and inflammatory responses, in both diseases [75, 76]. In addition, when compared with other diseases or tumor models, few common differential proteins overlapped with those associated with MDD, indicating that the urine proteome has the potential to distinguish different types of diseases. The biological processes associated with enriched differential proteins in different diseases were also compared with those associated with the CUMS model. As a result, some common biological processes were found, such as cell adhesion, the acute-phase response, complement activation and the positive regulation of gene expression were enriched. We therefore suggest that some specific biological processes related to MDD deserve more attention, including the negative regulation of nitric oxide mediated signal transduction and central nervous system development.

Overall, our results indicated that the CUMS model is a valuable model for the study of MDD. Through various unpredictable and mild stress stimulations of mice, the CUMS model leads to depressive-like behaviors, which simulates the stressful life of modern people. In addition, the urine proteome can also reflect the behavioral changes due to lacking pleasure. A total of 45 differential proteins were selected, 24 of which have been reported to be associated with MDD pathological mechanisms, while 10 have been suggested as MDD biomarkers. The effect of the growth and development in this research cannot be avoided. To further illustrate the application of our results, urine samples from MDD patients will be needed in future clinical studies.

## Acknowledgments

This work was supported by the National Key Research and Development Program of China (2018YFC0910202 and 2016YFC1306300), the Fundamental Research Funds for the Central Universities (2020KJZX002), the Beijing Natural Science Foundation (7172076), the Beijing Cooperative Construction Project (110651103), the Beijing Normal University (11100704), and the Peking Union Medical College Hospital (2016-2.27).

## Supporting Information

**Table S1.** Variable isolation window information for the DIA method with 39 windows.

**Table S2.** Number of proteins identified in each sample.

**Table S3.** Details of the quantification of each sample using the Spectronaut Pulsar X.

**Table S4.** Results of the random grouping test.

**Table S5.** Functional enrichment analysis results of differential proteins including biological processes, cellular components, molecular functions and pathways.

**Table S6.** Results of the comparation between the CUMS model and 15 other disease models, including urinary proteins and biological processes.

**Figure S1.**
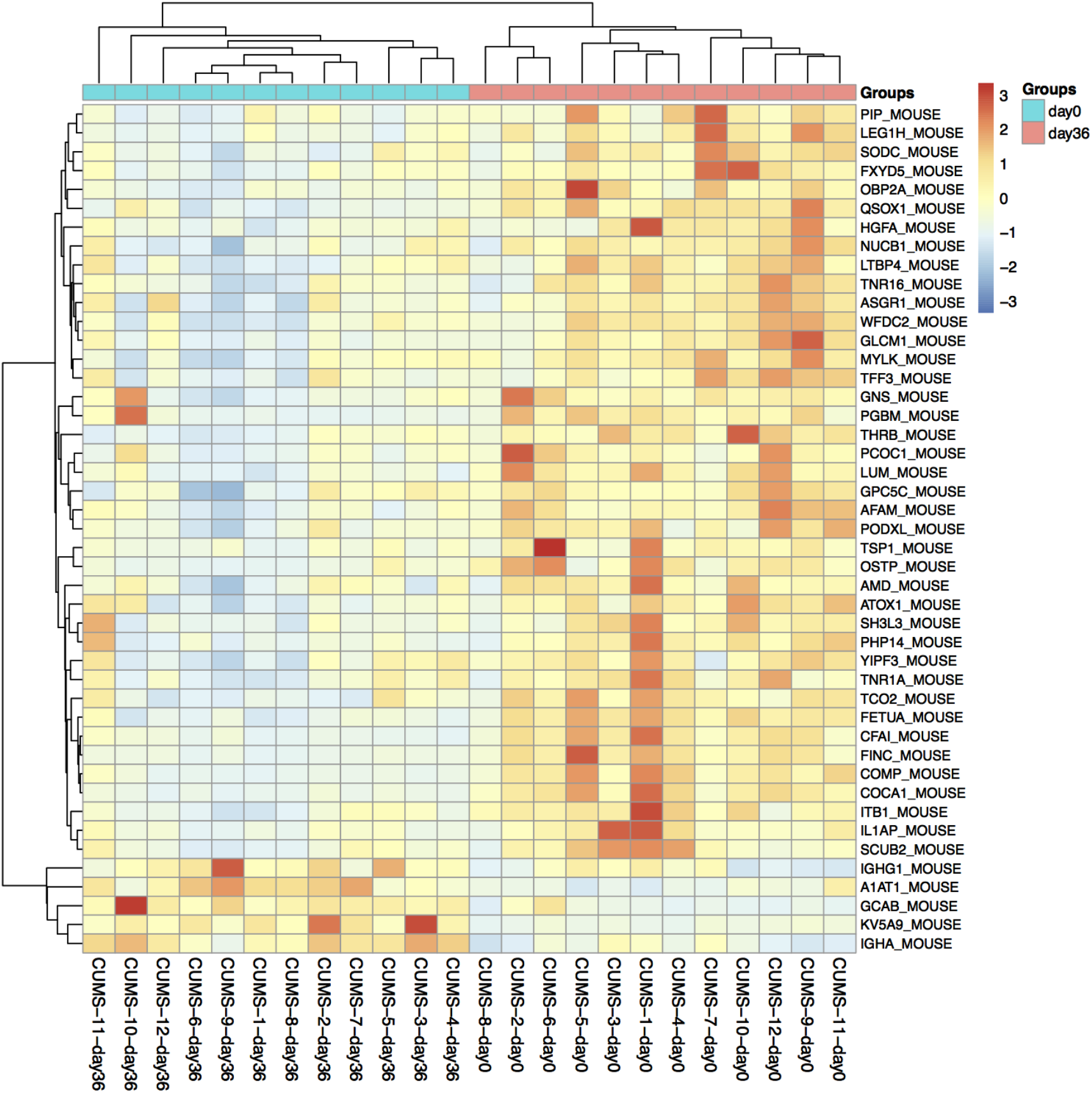
Hierarchical cluster analysis of 45 differential proteins.

